# Fabrication and electromechanical characterization of free-standing asymmetric membranes

**DOI:** 10.1101/2020.10.08.331470

**Authors:** Paige Liu, Oscar Zabala-Ferrera, Peter J. Beltramo

## Abstract

All biological cell membranes maintain an electric transmembrane potential of around 100 mV, due in part to an asymmetric distribution of charged phospholipids across the membrane. This asymmetry is crucial to cell health and physiological processes such as intracell signaling, receptor-mediated endocytosis, and membrane protein function. Experimental artificial membrane systems incorporate essential cell membrane structures, such as the phospholipid bilayer, in a controllable manner where specific properties and processes can be isolated and examined. Here, we describe a new approach to fabricate and characterize planar, free-standing, asymmetric membranes and use it to examine the effect of headgroup charge on membrane stiffness. The approach relies on a thin film balance used to form a freestanding membrane by adsorbing aqueous phase lipid vesicles to an oil-water interface and subsequently thinning the oil to form a bilayer. We validate this lipid-in-aqueous approach by analyzing the thickness and compressibility of symmetric membranes with varying zwitterionic DOPC and anionic DOPG content as compared to previous lipid-in-oil methods. We find that as the concentration of DOPG increases, membranes become thicker and stiffer. Asymmetric membranes are fabricated by controlling the lipid vesicle composition in the aqueous reservoirs on either side of the oil. Membrane compositional asymmetry is qualitatively demonstrated using a fluorescence quenching assay and quantitatively characterized through voltage-dependent capacitance measurements. Stable asymmetric membranes with DOPC on one side and DOPC/DOPG mixtures on the other were created with transmembrane potentials ranging from 15 to 80 mV. Introducing membrane charge asymmetry decreases both the thickness and stiffness in comparison to symmetric membranes with the same overall phospholipid composition. These initial successes demonstrate a viable pathway to quantitatively characterize asymmetric bilayers that can be extended to accommodate more complex membranes and membrane processes in the future.

**SIGNIFICANCE**
A defining characteristic of the cell membrane is asymmetry in phospholipid composition between the interior and exterior bilayer leaflet. Although several methods have been used to artificially create membranes with asymmetry, there has not been extensive characterization of the impact of asymmetry on membrane material properties. Here, a technique to fabricate free-standing asymmetric membranes is developed which facilitates the visualization and electromechanical characterization of the bilayer. Asymmetry in anionic phospholipid concentration is quantified by measurements of membrane capacitance at varying voltages, which also allows for determination of the membrane compressibility. This method represents an advance in the development of artificial biomembranes by reliably creating phospholipid bilayers with asymmetry and facilitates the interrogation of more complex biological processes in the future.

## INTRODUCTION

As the barrier between the cell’s interior and its outside environment, the cell membrane is a key interface in biological systems. Consequently, it has been a high priority for many years to understand the components of the cell membrane, and in particular to decipher the role of the phospholipid bilayer in myriad biological processes occuring in the cell. (1–5). In order to better understand the contribution of membrane structure and composition on the numerous functions the cell membrane performs, an array of artificial biomembrane systems have been employed to control experimental parameters in order to extract quantitative understanding of membrane material properties. Systems that are widely in use include vesicle based techniques such as liposomes and giant unilamellar vesicles (GUVs)(6, 7) and planar systems such as droplet interface bilayers (DIBs)(8–10), supported lipid bilayers (SLBs)(11, 12), and black lipid membranes (BLMs)(13, 14). These systems have been used to conduct numerous studies characterizing the biophysical properties of lipid bilayers, such as the determination of membrane bending rigidity via micropipette aspiration (15, 16), optical flicker spectroscopy (17, 18) or small-angle x-ray scatting (19, 20), as well as the electromechanical properties of DIBs (21) or BLMs (22–24). The majority of this work has been performed using symmetric membranes, where each leaflet has the same phospholipid composition.

However, the cell membrane possesses an asymmetric phospholipid composition where, in particular, the interior leaflet of the cell is enriched with negatively charged phospholipids compared to the exterior leaflet (25–30). This membrane charge distribution drives interaction with macromolecules at interface (31), dictates biological processes such as vesicle budding (32), controls the conformation of membrane proteins (33, 34), and becomes altered when cells undergo cancerous transformation (35–42). As a result, there has been a recent surge of interest in fabricating asymmetric model systems for the systematic study of the role of phospholipid asymmetry in biological systems. The techniques described above have all recently been extended to asymmetric membranes. For example, asymmetric vesicles and GUVs can be formed using cyclodextrin-mediated lipid exchange (43, 44), from inverted emulsion transfer (45, 46), or by using a jet-flow method (47). Asymmetric SLBs formed from vesicle adsorption(48), Langmuir-Blodgett techniques (49), and cyclodextrin-mediated exchange (50) have shown interesting domain coupling between leaflets. The DIB approach to creating membranes can readily accomodate leaflet asymmetry (51, 52), and free-standing asymmetric BLMs may also be formed using the Montal-Mueller technique(53). However, there remains a dearth of systematic studies quantifying the relationship between compositional asymmetry and membrane material properties, with the notable exception of recent GUV experiments that determined that the bending moduli of asymmetric membranes was 50% greater than their symmetric counterparts (54, 55).

We have recently developed a microfluidic platform to create planar, freestanding, large area model biomembranes (LAMBs) which allows for characterization of membrane elasticity and compressibility(56–58). In this work, we extend the LAMB platform to fabricate asymmetric phospholipid membranes and focus on the effect of headgroup charge on the membrane Young’s modulus. Previously, only symmetric membranes were able to be formed by this method because it relied on lipids suspended in an oil solvent (lipid-in-oil). To create membrane asymmetry, we first establish the use of lipids suspended in independent aqueous buffers on either side of the membrane (lipid-in-aqueous). We validate the lipid-in-aqueous approach by comparison to the lipid-in-oil method using symmetric membranes formed from binary mixtures of zwitterionic and anionic phospholipids. For the asymmetric studies, we start by qualitatively demonstrating the fabrication of an asymmetric bilayer using a fluorescence quenching assay. We then quantitatively characterize the offset potential and increase in bilayer compressibility that occurs upon introducing membrane charge asymmetry. The results demonstrate the ability of this platform to quantitatively characterize asymmetric membranes using this new fabrication approach.

## MATERIALS AND METHODS

### Materials

The phospholipids 1,2-dioleoyl-sn-glycero-3-phosphocholine (DOPC), 1,2-dioleoyl-sn-glycero-3-phospho-(1’-rac-glycerol) sodium salt (DOPG), and 1,2-dioleoyl-sn-glycero-3-phosphoethanolamine-N-(lissamine rhodamine B sulfonyl) ammonium salt (DOPE-Rh) were obtained from Avanti Polar Lipids. Sodium chloride (NaCl), calcium chloride (CaCl_2_), sodium bicarbonate (NaHCO_3_), hexadecane and octadecyltrichlorosilane (OTS) were purchased from Fisher Scientific. All aqueous solutions are prepared using ultra-pure water (Milli-Q, Millipore-Sigma). Before use, hexadecane is filtered through an 0.2 *µ*m aluminum oxide mesh. QSY-7 succinimidyl ester was obtained from Life Technologies Corporation. A custom glass microfluidic chip with a ‘bikewheel’ geometry used for the LAMB platform is ordered from Micronit Microfluidics. Full details of the device design, geometry, and operation can be found in prior work(56). Briefly, the microfluidic chip contains a series of channels in a bikewheel and spoke geometry which allow for the uniform lateral drainage of solution from a 0.9 mm diameter hole drilled in the center of the chip. Before use, the microfluidic chip is hydrophobized in a 1 *µ*M solution of OTS in hexadecane, then plasma cleaned for 5 seconds to partially remove the hydrophobic coating from the exterior of the chip.

### Bilayer Fabrication

The method to prepare bilayers from a lipid-in-oil suspension was previously discussed by Beltramo et al. (56), here we focus on the relevant experimental details for the current work creating asymmetric membranes from a lipid-in-aqueous solution. First, the teflon sample holder for the microfluidic chip is modified to allow for solution exchange and compartmentalization between the aqueous phase above and below the phospholipid membrane Figure 1A. The microfluidic chip is installed in the sample holder using vacuum grease to create a seal between the top (orange) and bottom (blue) aqueous chambers, allowing for both chemical and electrical isolation of the chambers. The microfluidic chip is connected to a capillary tube which allows for control of the capillary pressure within the central orifice using a syringe pump (Harvard Apparatus PHD Ultra) and pressure transducer (MKS PR 4000B-F). The sample chamber is designed such that channels connect the two side chambers to the area underneath the microchip (blue regions in Figure 1A). In addition, the two chambers are connected with PTFE tubing to a push-pull syringe pump. These two details allow for equal hydrostatic pressure on the bilayer, and the exchange of aqueous solution conditions underneath the microchip.

**Figure 1:**
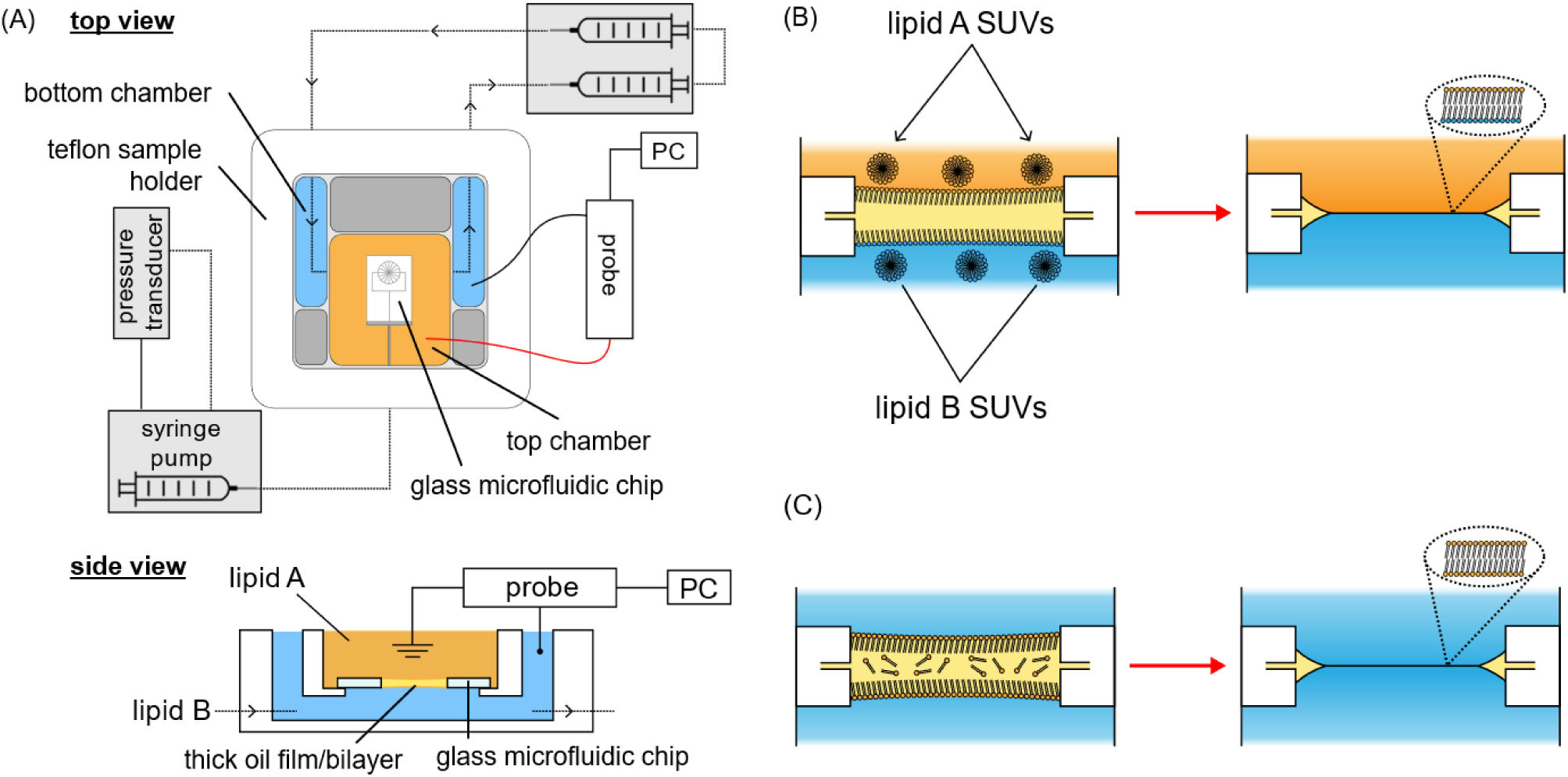
(A) Schematic of the platform used to form asymmetric bilayer membranes. The lipid bilayer is formed in the center of the glass microfluidic chip, which is electrically and chemically isolated between the top (orange) and bottom (blue) aqueous chambers. A push-pull syringe pump allows for solution exchange to introduce lipid B below the microfluidic chip, while pipetting is used to introduce lipid A in the top chamber. Once the lipids adsorb to the hexadecane-aqueous interface, the oil in the microfluidic chip is controllably thinned using a syringe pump and pressure transducer, resulting in an asymmetric bilayer (B). Alternatively, the microchip can be filled with a lipid-in-oil solution to form symmetric bilayers (C). Not to scale.

To form asymmetric membranes, two different aqueous solutions with the desired phospholipid compositions (lipid A and lipid B) are prepared. Lipid mixtures are prepared using stock dilutions in chloroform and dried in scintillation vials under nitrogen. The vials are then placed under high vacuum (≤5mbar) overnight to ensure the removal of any residual chloroform.

The dried lipid is then resuspended in an aqueous buffer (150 mM NaCl, 2 mM CaCl, and 0.2 mM NaHCO_3_, filtered through an 0.2 *µ*mm pore filter prior to use) at a concentration of 0.1 wt%. This aqueous solution is then sonicated in a water bath for 6-8 hours to convert the lipids into small unilamellar vesicles (SUVs). To create a bilayer, the microfluidic chip is filled with hexadecane and loaded into the sample chamber with aqueous buffer (without lipid) on either side. A thick oil film is formed to seal off the top and bottom chambers, and subsequently the chambers are filled with lipid A and lipid B using pipette exchange or the push-pull syringe pump system, respectively. With the aqueous lipid solution on either side, the thick oil film of hexadecane is left for at least 30 minutes to allow lipids to adsorb to the oil-water interface. Finally, an asymmetric bilayer is formed by thinning the oil film (Figure 1B). Symmetric bilayers are readily formed from aqueous phase lipids using this approach, without having to go through the aqueous solution replacement steps. Symmetric bilayers can also be formed from resuspending dried lipids directly in hexadecane at a concentration of 2.5 mg/mL, as done previously (56). The lipid-in-oil solution is then sonicated in a water bath for at least 2 hours before use. The microfluidic chip is filled with the lipid-in-oil solution, loaded into the chamber with aqueous buffer on either side, and upon subsequent thinning of a thick oil film symmetric bilayers are formed (Figure 1C).

### Material Property Characterization

To characterize the thickness and stiffness of a bilayer, as well as measure the offset potential of charge asymmetric bilayers, a patch clamp amplifier system (HEKA) is used with a pulse generator file (PGF) to automate the voltage application and data collection process. Alternating positive and negative voltages are applied in 1 second “on” pulses at an amplitude ranging from 25 mV to 200 mV with a 2 second “off” pulse at 1mV in between. The capacitance across the bilayer during these voltage pulses is recorded. At the same time, the bilayer is imaged at 10 FPS, and the changing area of the bilayer during the experiment is determined using a custom MATLAB script. An example data set resulting from this experiment is shown in Figure 2. The capacitance (*C*) and area (*A*) data is used to calculate the voltage-dependent hydrophobic thickness of the bilayer by *d* = *εε*_0_ *A*/*C*, where *ε*_0_ is the permittivity of free space. We use 2.5 as the dielectric constant of the bilayer, *ε*, in accordance with prior work (57). The degree to which the membrane thins as a result of the applied voltage is directly related to the membrane Young’s modulus (compressibility perpendicular to the plane of the membrane) by (23, 24, 57):

**Figure 2:**
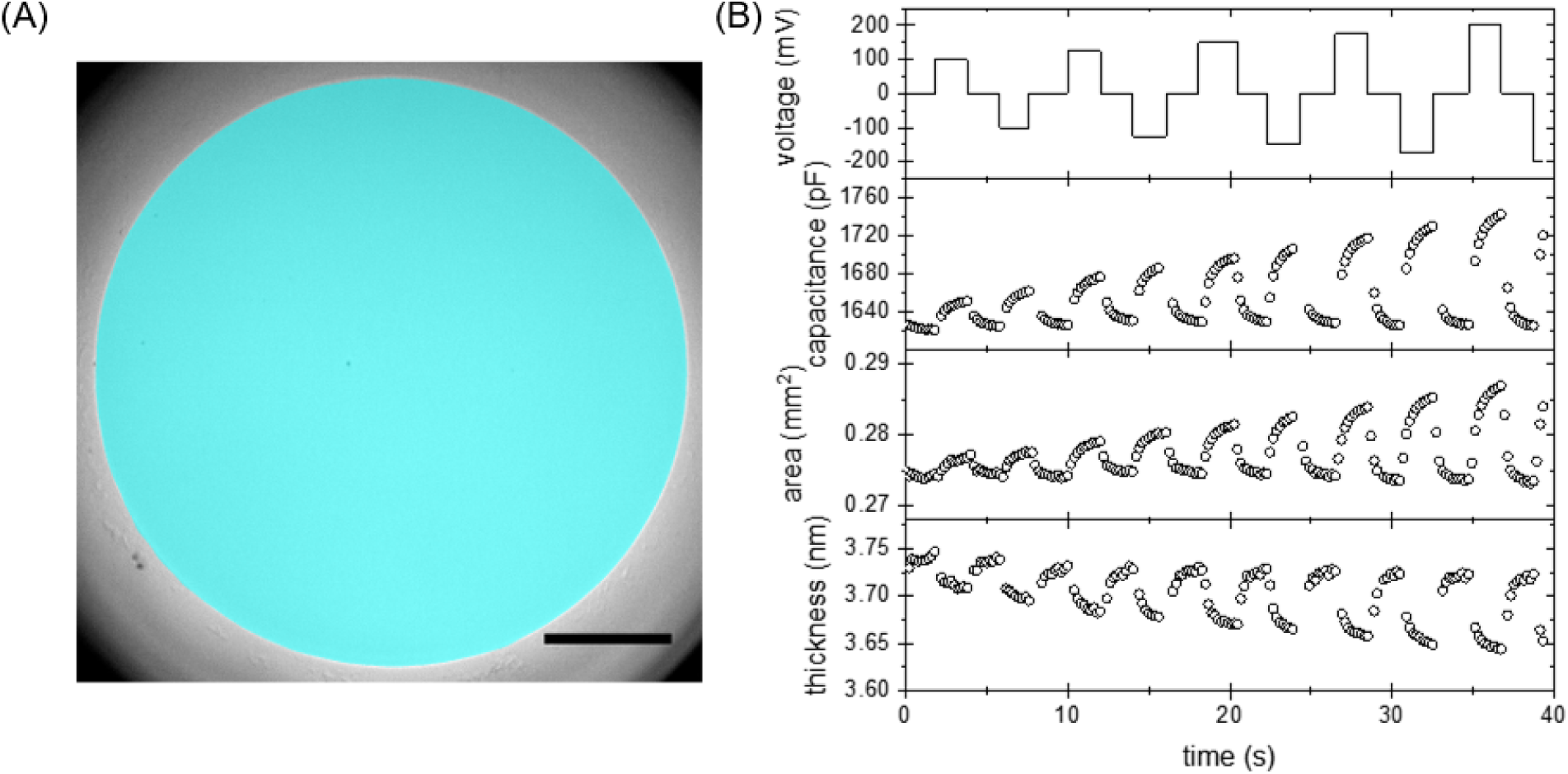
(A) Image of a formed bilayer with the membrane area detected by image processing false colored in blue. Scale bar is 100 *µ*m. (B) Exemplary experiment applying a voltage sweep while measuring membrane area, capacitance, and thickness.

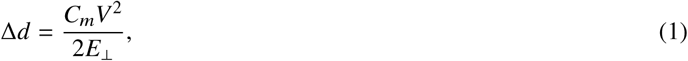

where Δ*d* is the decrease in thickness from the static thickness, *C*_*m*_ is the static membrane specific capacitance, *V* is the applied voltage, and *E*_⊥_ is the Young’s modulus. We find that as the applied voltage approaches zero there is increased noise in the capacitance data, therefore to determine the static specific capacitance and hydrophobic thickness of symmetric membranes we fit a parabola to all the voltage-thickness data for a given bilayer and report the static thickness as the fitted value at *V* = 0. This often corresponds well with the thickness measured during our experiments using *V* = 1mV. For asymmetric membranes, the parabolic fit also determines the offset voltage where the thickness of the membrane is at its maximum, as explained below.

### Fluorescence Quenching Assay

To demonstrate the formation of an asymmetric bilayer, a fluorescence assay was conducted by adding QSY-7, a black hole amine-based quencher, to membranes formed with 1 mol% DOPE-Rh. Three assays were run-introducing the QSY-7 quencher to one side of a fluorescently symmetric bilayer and the two sides of an asymmetric bilayer with fluorescent headgroup lipid only on one side. The fluorescence was monitored before and after quench and measured using ImageJ in ten separate areas on each bilayer image to obtain ten fluorescent intensities. These intensities were then averaged and scaled by the pre-quench intensities to obtain scaled fluorescent intensities before and after quench.

## RESULTS AND DISCUSSION

### Comparison between Lipid-in-oil and Lipid-in-aqueous Symmetric Membranes

Previous work performed with the LAMB platform created bilayers from lipids suspended in an oil phase, which was effective in controlling the phospholipid composition in symmetric bilayers (57) but could not produce asymmetric membranes since the lipids would adsorb to both sides of the oil-aqueous interface. To generate membrane asymmetry with this platform, it is necessary to isolate the aqueous compartments on either side of the oil interface and introduce phospholipids from the aqueous phase. We first validate this fabrication approach by analyzing the thickness and compressibility of symmetric membranes with the same composition formed from either the original lipid-in-oil approach or the new lipid-in-aqueous method using binary DOPC/DOPG bilayers. Visually, there is no difference in bilayer formation in either method-after the thick oil film thins and interferometric fringes appear the nucleation and growth of a circular black lipid membrane follows, whose area can be controlled by the capillary pressure in the microfluidic chip. We do find that it is necessary to ensure that there is an adequate concentration of lipids at the oil-aqueous interface before thinning for successful bilayer formation to occur. With lipids suspended in oil at a concentration of 0.3 −0.6 wt%, this happens relatively rapidly and the bilayer may be formed almost immediately after creating the thick oil film. However, with lipids suspended in the aqueous phase at a lower concentration of 0.05 wt% to 0.2 wt%, it is necessary to wait at least 30 minutes to ensure an adequate interfacial concentration, or a bilayer will not nucleate upon thinning. An analogous factor to bilayer formation has been found using droplet interface bilayers (59), where the droplet oil-water interface needed to be compressed to more tightly pack the DOPC monolayer and increase the probability of successful bilayer formation. In our set up, thinning the oil from a thick film may compress each monolayer in a similar fashion. However, empirically we have found that immediately thinning the oil film after adding lipids to the aqueous phase results in the brief nucleation and rapid bursting of the thin film, indicating that there also must be adequate time for lipids to adsorb to the interface in addition to the slight compression upon thinning. After formation, the bilayers created here have membrane tensions in the range of 1 −5 mN/m, can withstand voltage potentials as high as 200mV, and can exist for over 2 hours under repeated compression, indicating they are fully packed and stable bilayers.

Once formed, membranes are characterized using the voltage pulse method described above. By lining up capacitance measurements with area measurements derived from simultaneous imaging, we calculate the specific capacitance and thickness for each applied voltage, as shown in Figure 3A-B. As expected for symmetric membranes (24), the specific capacitance and thickness of the membranes are quadratic with respect to voltage and centered at zero. Since the slope of the data in Figure 3C is inversely related to the membrane Young’s modulus per Equation 1, it is obvious that increasing headgroup charge concentration increases the stiffness of symmetric membranes.

**Figure 3:**
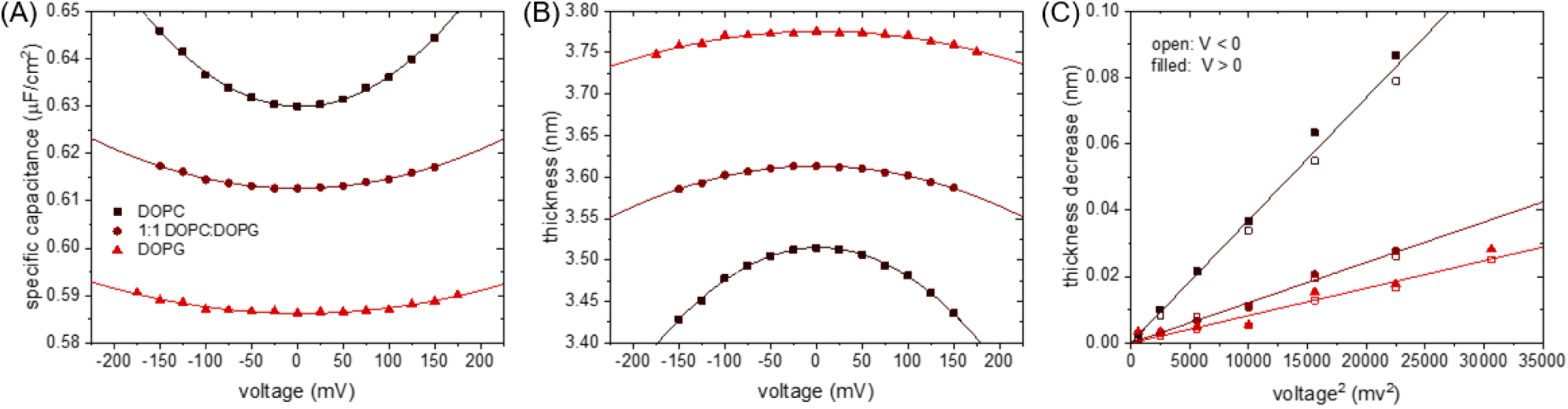
(A) Specific capacitance and (B) thickness for symmetric lipid membranes formed from the lipid-in-aqueous approach with varying DOPC/DOPG content. The bilayer compresses symmetrically with voltage, as shown in (C). The membrane Young’s modulus is inversely related to the slope of the thickness vs. V^2^ relationship, increasing with DOPG content.

The compiled data for DOPC/DOPG bilayers formed by either the lipid-in-oil or lipid-in-aqueous approach is shown in Figure 4. We found minimal dependence of static thickness with DOPG content in a DOPC-DOPG binary mixture using the lipid-in-oil approach, however, the hydrophobic thickness consistently increased when adsorbing lipids from the aqueous phase (Figure 4A). It has previously been shown that membranes formed from hexadecane have larger thickness than their solvent-free counterparts due to residual oil molecules remaining in the membrane(21, 57), which has been estimated to be ∼36% using molecular dynamics simulations (60). The increase in thickness with DOPG concentration could be due to anionic DOPG introducing increased electrostatic repulsion between headgroups on opposing leaflets, however existing literature measuring the effect of headgroup charge on hydrophobic bilayer thickness is mixed (61–63). Temperature-dependent x-ray diffuse scattering of oriented stacks and unilamellar vesicles of DOPC (61) indicate a consistently smaller thickness (2.68 nm at 30 ^°^C) than a combination of small-angle x-ray and neutron scattering (SAXS/SANS) of unilamellar DOPG vesicles (2.82 nm at 30 ^°^C)(62). However, differences in measurement technique, lipid preparation, and experimental error may all contribute to this thickness difference. The hydrophobic thickness of DPPG (62) and DOPC (63) unilamellar vesicles determined by SANS/SAXS at 50 ^°^C are within experimental error (2.78 nm for DPPG compared to 2.85 nm for DOPC). This agrees with data in the same papers comparing the thickness of POPC (63) and POPG (62) which also shows a slight thickness decrease with anionic headgroup bilayers. In any case, the hydrophobic thickness of a lipid bilayer membrane is principally determined by the acyl chain length and saturation. The increase in membrane thickness measured in the current work compared to scattering experiments on solvent free bilayers can be attributed to the prescence of residual oil molecules in the bilayer, and the choice of dielectric constant further hinders quantitative comparisons between thicknesses determined by capacitance and scattering measurements. The dielectric constant of the hydrophobic phase is taken as 2.5, consistent with our prior work(57). Others have used dielectric constants as low as 2.2 for DPhPC lipid bilayers (64), which would decrease the capacitive thickness reported here. If the ∼36% by volume residual hexadecane in the bilayer, as suggested by molecular dynamics simulations(60), also increases the hydrophobic thickness by the same amount, the DOPC x-ray scattering thickness is roughly in line with our results, giving further confidence to our choice of dielectric constant. Residual oil, combined with experimental uncertainty, may also help explain the greater dependence of thickness on headgroup chemistry shown in Figure 4A.

**Figure 4:**
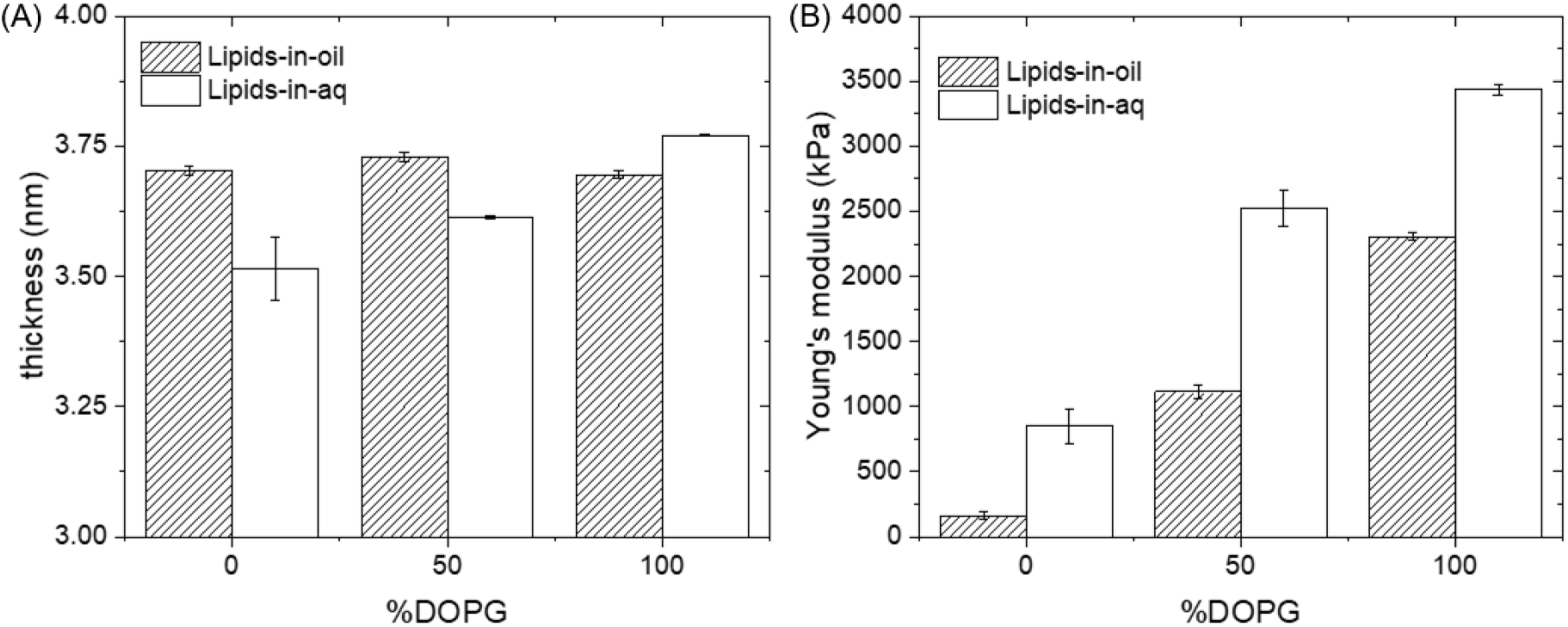
(A) The dependence of hydrophobic membrane thickness on DOPC/DOPG concentration and the method of bilayer fabrication. (B) Increasing DOPG content stiffens membranes, while membranes formed from the aqueous phase are more stiff than their oil counterparts.

The Young’s modulus was found to increase with increasing DOPG content regardless of fabrication method (Figure 4B).As the concentration of anionic lipids in the membrane increases, one would expect the membranes to become monotonically stiffer and less compressible as seen here. When bilayers of the same composition were fabricated using the lipid-in-aqueous method, they were found to have higher moduli. Our results are consistent with bending modulus measurements of GUVs with varying DOPG concentration performed using optical tweezers which exhibited the same trend of increasing modulus with higher DOPG content (65).

### Asymmetric Membranes

To qualitatively demonstrate the formation of asymmetric membranes with independent aqueous solution conditions on either side of the bilayer, a series of quenching assays were performed. As a control, a symmetric bilayer composed of DOPC and 1 mol% DOPE-Rh (DOPC:DOPE-Rh 99:1) was formed using the lipid-in-oil approach before introducing QSY-7 to the top compartment of the bilayer. Satisfactorily, the fluorescence intensity was found to decrease to about 55% of the starting intensity with minimal variation in fluorescence intensity within the bilayer area, indicating that one leaflet was fully quenched (Figure 5A).

**Figure 5:**
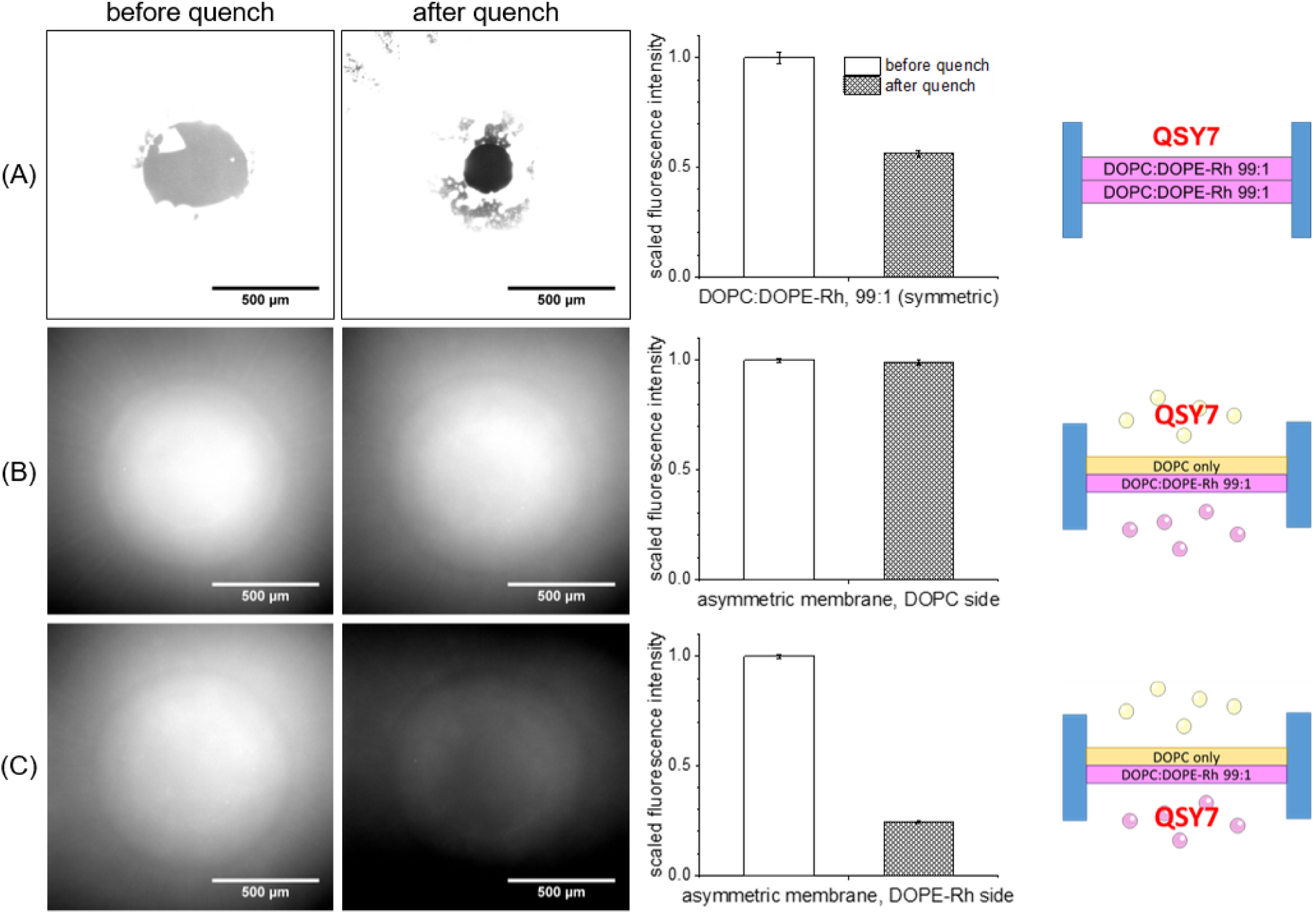
(A) Fluorescence quenching of a symmetric DOPC membrane formed from the oil phase using QSY-7 added to one side shows the chemical isolation between the two sides of the membrane and a ∼50% decrease in fluorescence intensity, indicating one leaflet is fully quenched. Fluorescence quenching of an asymmetric membrane with fluorescence headgroup on only one side of the membrane shows (B) no decrease in intensity when the quencher is added to the opposite side and (C) a steep decrease in intensity when the quencher is added to the same side of the fluorescent headgroup phospholipid.

An asymmetric bilayer was then formed using the lipid-in-aqueous approach with DOPC vesicles on one side and DOPC:DOPE-Rh 99:1 vesicles on the other. When QSY-7 was added to the side containing no DOPE-Rh, there was a negligible change in fluorescence intensity (Figure 5B). However, when QSY-7 was added to the bottom of the asymmetric bilayer, which was the leaflet composed of DOPC:DOPE-Rh 99:1, there was about an 80% drop in fluorescence intensity, as shown in Figure 5C. The remaining fluorescence measured may be explained by background noise generated by unquenched vesicles in the aqueous solution away from the bilayer.

The fluorescence experiments done to validate the lipid-in-aqueous method of generating asymmetric planar membranes bring about two conclusions. One, it is clear that the oil film separates the two components of aqueous lipids so that the asymmetric membrane can form without lipid mixing. Secondly, once the membrane is formed we do not observe lipid flip-flop within the duration of a typical experiment (∼three hours). This is supported by experiments tracking lipid flip-flop in SLBs or GUVs, which have found this process takes at least half a day to occur (66). While bilayer asymmetry is not a thermodynamically stable state, there is still a high energy cost associated with the lipid flip-flop mechanism and it is considered a metastable equilibrium state (67). From the data shown, there is not strong evidence that lipid flip-flop has occurred within the experimental time frame, so we believe that the asymmetry we generate is stable enough for characterization experiments described below.

In an asymmetrically charged bilayer, the membrane holds a charge in the absence of an applied voltage (30). This is opposed to an asymmetric bilayer with solely zwitterionic phospholipids, and a symmetric bilayer with or without charged lipids, which will all possess a net zero charge in the absence of any applied voltage. Normally when some voltage is applied to the bilayer, it will be compressed. However, in the case of an asymmetrically charged bilayer, if the applied voltage is equal and opposite to the membrane charge, the membrane will become unstressed, relax, and increase in thickness. Experimentally, this is seen in the shift of the vertex of the quadratic fit of the thickness to voltage data. In an asymmetric bilayer created with one leaflet of 100% DOPC and one leaflet of 50% DOPC and 50% DOPG (notation: DOPC/(DOPC:DOPG 1:1)), this offset voltage was measured to be 15 mV, which can be seen in Figure 6A. This shift must be taken into account in determining the membrane

**Figure 6:**
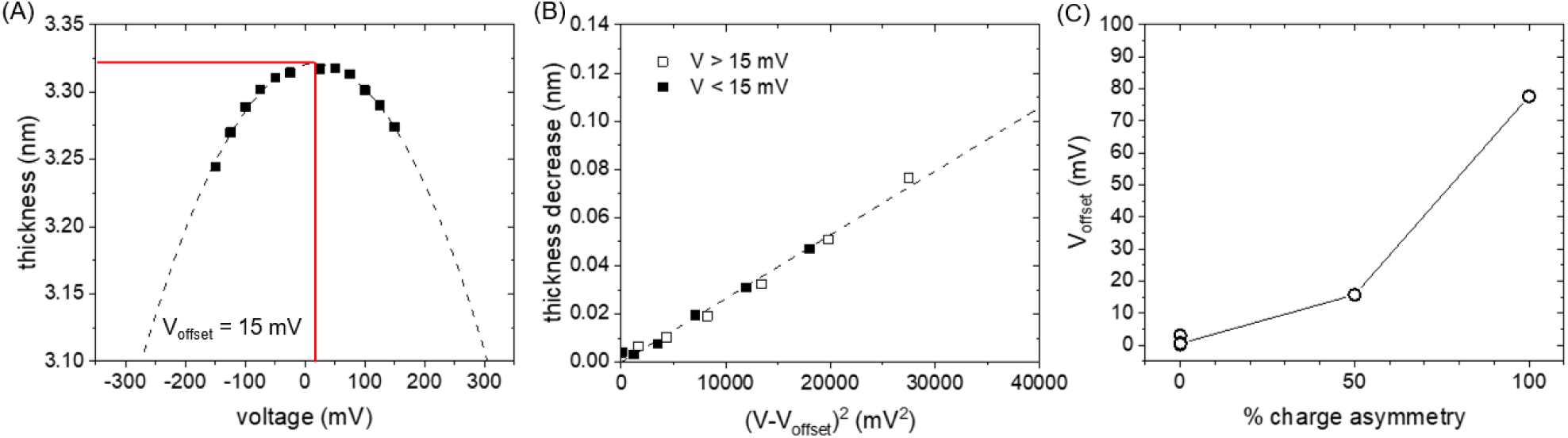
(A) Thickness of an asymmetric DOPC/(DOPC:DOPG 1:1) bilayer shows asymmetric compression upon voltage application. (B) By considering the offset in potential caused by the charge asymmetry, the data still follow the expected V^2^ dependence. (C) The offset potential increases with the degree of membrane charge asymmetry, as defined in the text.

Young’s modulus, such that

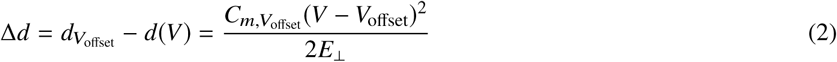

where 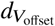 and 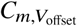 is the thickness and specific capacitance, respectively, at the offset voltage, *V*_offset_. This shift is shown in Figure 6B, where the thickness decrease from the unstressed state is identical on either side of the offset voltage. After taking the offset voltage into account, the thickness data once again follow the expected voltage squared relationship, which allows for determination of the Young’s modulus of asymmetric membranes. We have characterized bilayers with varying degrees of charge asymmetry, defined as

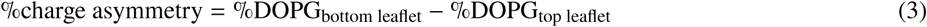

Therefore, the aforementioned DOPC/(DOPC:DOPG 1:1) bilayer contains 50% charge asymmetry while a fully asymmetric bilayer with DOPC on one leaflet and DOPG on the other contains 100% charge asymmetry. As the charge asymmetry increases, so does the membrane offset potential, as shown in Figure 6C. For a membrane with 100% charge asymmetry, the offset potential approaches 80 mV. This is close but still lower than the 100 mV offset potential commonly found in cells(68), whose resting potential is due to osmotic pressure, curvature, and charged membrane proteins in addition to phospholipid charge asymmetry.

In order to deduce the effect of charge asymmetry on the material properties of the membrane, it is necessary to compare membranes with the same overall membrane composition but with different leaflet compositions. This is shown pictorially in Figure 7A, where we compare symmetric membranes with 25% and 50% DOPG with their maximally asymmetric counterparts and symmettric membranes composed of only one phospholipid. It is important to reiterate that all of these membranes possess the same acyl chain content and only vary in headgroup chemistry. Introducing headgroup charge asymmetry decreases the unstressed membrane thickness and also decreases the membrane Young’s modulus (Figure 7B,C). Both of these observations are consistent with the decreased electrostatic repulsion between leaflets that occurs in the asymmetric case. Symmetric membranes with DOPG on both leaflets are expected to have electrostatic repulsion that scales with the DOPG concentration, which causes the membranes to become thicker and stiffer. When this is removed the membranes are thinner and compress more under an electric stress. The only difference between the two asymmetric membranes measured is in the lateral headgroup interactions on the leaflet that has varying DOPG content since the other leaflet is 100% DOPC. We do not see meanginful differences between the thickness of the two asymmetric membranes, however there are hints that increasing charge asymmetry increases the Young’s modulus of the membrane. This is difficult to resolve conclusively in the current work, but it appears to be secondary compared to the stark effects of asymmetry.

**Figure 7:**
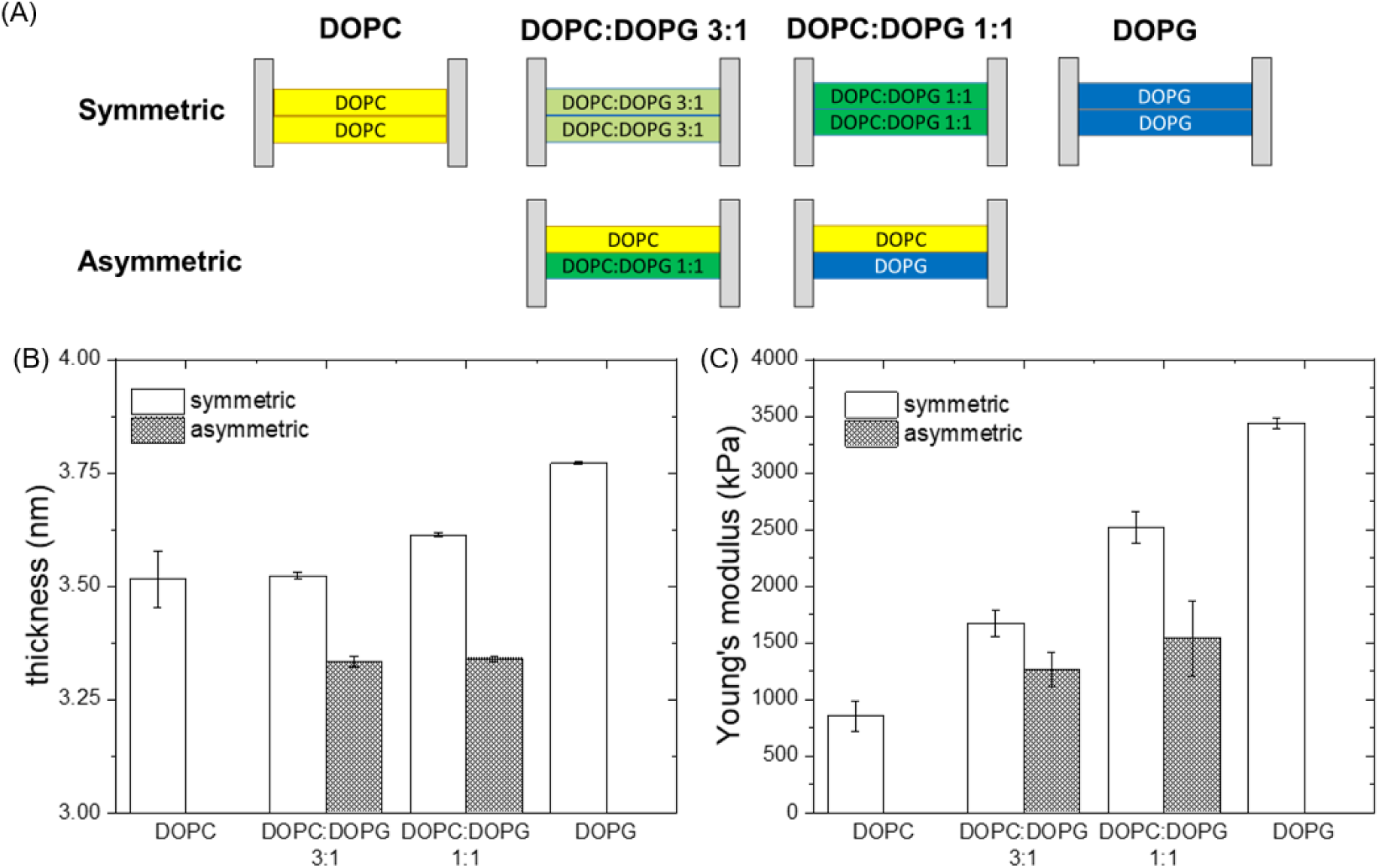
(A) Pictogram of the four symmetric and two asymmetric bilayer compositions studied. The unstressed hydrophobic membrane thickness (B) and the membrane Young’s modulus for all six membranes formed from the lipid-in-aqueous approach.

It is worthwhile to consider our results in light of recent experimental (54, 55) and theoretical studies(67) which found an increase in membrane bending rigidity with increasing leaflet asymmetry. In contrast to our work, both prior results were determined using membranes with asymmetry introduced through varying acyl chain lengths/saturation on phospholipids with zwitterionic headgroups and reported bending rigidities, not compressibility. While we previously discussed the decrease in thickness and Young’s modulus with variable charge asymmetry, one can look at a subset of the data here with DOPC on one leaflet and varying DOPC/DOPG content on the opposing leaflet (DOPC, DOPC:DOPG 1:1, DOPG). These three membranes lack the increased electrostatic repulsion present in the three symmetric membranes with DOPG in both leaflets, however as compositional asymmetry increases the membranes stiffen similar to what was found by others. One possible explanation for this is the development of a differential stress across the two leaflets(67), especially since the results are not merely a compositional weighting of the Young’s modulus found on symmetric bilayers. We also note that the fully asymmetric DOPC/DOPG membrane is less stiff than the average stiffness of the 100% DOPC and 100% DOPG. This is also the case for the asymmetric DOPC/(DOPC:DOPG 1:1) membrane. Both interactions orthogonal and parallel to the plane of the membrane contribute to the stiffness changes of charge asymmetric membranes, and to fully decouple each effect additional experiments with varying compositional, acyl tail, and headgroup asymmetry will need to be studied in the future. In total, these results demonstrate a novel improvement on the fabrication and characterization of free-standing asymmetric membranes with independent compositional control over both leaflets of the membrane.

## CONCLUSION

The LAMB platform has been used to systematically characterize the material properties of symmetric bilayers with varying anionic phospholipid headgroup concentrations formed from the previous lipid-in-oil approach and the new lipid-in-aqueous method. The lipid-in-aqueous method further allows for the fabrication of freestanding membranes with leaflet asymmetry. The effect of charged headgroups in a lipid bilayer has been quantified in terms of its effect on static thickness, Young’s modulus, and offset voltage. Increasing the concentration of charged DOPG phospholipids in symmetric bilayers formed using the lipid-in-aqueous approach causes an increase in the static thickness and modulus of the bilaye. Stable asymmetric bilayers were formed and verified qualitatively using a fluorescence quenching assay, and quantitatively verified by the detection of an offset voltage from capacitance measurements of a charge asymmetric membrane. As expected, increasing the amount of membrane charge asymmetry increases the membrane offset potential. When comparing membranes formed with the same overall phospholipid composition, bilayers with charge asymmetry have decreased thickness and Young’s modulus in comparison with their symmetric counterparts, likely due to decreasing electrostatic repulsion between the opposing leaflets. Asymmetric bilayers with one leaflet composed of entirely zwitterionic phospholipids exhibit an increase in stiffness with increasing anionic lipid content in the opposing leaflet. These studies lay the groundwork for additional work involving more complex lipid mixtures with the goal to systematically measure the effects of perturbations of composition and solution conditions on membrane material properties.

## AUTHOR CONTRIBUTIONS

PJB designed the research and acquired funding for it. OZ performed and analyzed lipid-in-oil experiments. PL performed and analyzed lipid-in-aqueous experiments. All authors discussed the findings, experimental plans, and wrote the manuscript.

## ACKNOWLEDGMENTS

Funding for this research was provided by NSF under Award No. CBET-1942581 and the University of Massachusetts Amherst. The authors also acknowledge the UMass Institute for Applied Life Sciences AddFab facility for assistance fabricating sample cells used in this work.

